# DTH : A nonparametric test for homogeneity of multivariate dispersions

**DOI:** 10.1101/2025.06.29.662200

**Authors:** Asmita Roy, Glen A. Satten, Ni Zhao

## Abstract

Testing homogeneity across groups in multivariate data is an important scientific question in its own right, as well as well as an auxiliary step in verifying the assumptions of ANOVA. Existing methods either construct test statistics based on the distance of each observation from the group center, or as the mean of pairwise dissimilarities among observations in a group. Both approaches can fail when mean within-group distance is similar across groups but the distribution of the within-group distances are different. This is a pertinent question in high dimensional microbiome data, where outliers and overdispersion can distort the performance of a mean-dissimilarity-based test. We introduce the non-parametric Distance based Test for Homogeneity (`DTH`) which measures dissimilarity between groups by comparing the empirical distribution of within-group dissimilarities using a combination of the Kolmogorov-Smirnov and Wasserstein distances. For more than two groups, pairwise group tests are combined using a permutation-based p-value. Through simulations we show that our method has higher power than existing tests for homogeneity in certain situations and comparable power in other situations. We also provide a simple framework for extending the test to a continuous covariate.

## 1 Introduction

One approach to high-dimensional data analysis is to reduce the data to a matrix of pairwise dissimilarities. This approach is ubiquitous in analyses of data from microbiome studies, where ecologists have provided dissimilarity measures such as the Bray-Curtis, UniFrac or Aitchison dissimilarities that allow ecologically meaningful comparisons between samples. Unfortunately, reducing data in this way can make standard analyses more difficult. For example, regression analysis posits the effect of covariates at the individual level, not on pairs of individuals. Interestingly, there are circumstances in which it is still possible to conduct the standard analysis on the paired data. For example, if matrix of pairwise dissimilarities is Euclidean (i.e., has only non-negative eigenvalues), then `PERMANOVA`(Anderson [2001]) can be used to calculate F-tests for the effect of covariates in a regression setting. However, if the dissimilarity matrix is non-Euclidean (e.g., for the Bray-Curtis dissimilarity applied to count data), then these F-statistics can be negative. Another difficulty with reducing data to pairwise dissimilarities is that `PERMANOVA` is sensitive to both shifts in location and dispersion (scale) (Warton et al. [2012]). Thus, using `PERMANOVA` alone, it is difficult to tell if differences between groups, or along a covariate gradient, are differences in location, differences in dispersion, or some combination of each. Hence, it is of interest to develop tests of pure dispersion for pairwise dissimilarity data, to understand how groups may actually differ.

In the simplest case, where we compare two or more groups, and where the dissimilarity is Euclidean, standard multi-dimensional scaling techniques can be used to represent the data as points in a Euclidean space centered about a common zero. Then, the Euclidean distance between each point and the mean (or median) of its group can be evaluated directly using Levene’s test for equal variances(Brown and Forsythe [1974]). This is the approach taken by `PERMDISP` (Anderson [2006]) and implemented in the `betadisper` function in the `VEGAN` R package. This approach does not extend easily to the question of how dispersion varies along a continuous gradient. For non-Euclidean dissimilarities, Anderson proposes a somewhat heuristic modification to the distance to the mean (or median), but if this approach is used it is not completely clear what is being tested.

Gijbels and Omelka (Gijbels and Omelka [2013]) developed a test for dispersion for pairwise dissimilarity data that allows valid comparisons of within-group variances when the dissimilarity is non-Euclidean. Most notably, this approach does not require the data be representable in a Euclidean space, but instead computes the mean of all pairs of within-group distances. For a Euclidean matrix, it is known the sum of squared distances is equivalent to the mean of squared sums of distances from each observation to its group centroid, and also corresponds to the *U* -statistic form of the standard variance. They also give a version for use with the spatial median, but only consider grouped data, not continuous covariates. Importantly, they give a clever permutation scheme that allows generation of null replicates with equal dispersion that accounts for differences in location. Gijbels and Omelka propose two tests that we denote GO (calculates p-values using an asymptotic distribution) and GO.perm (calculates p-values by permutation).

Both Anderson and Gijbels and Omelka only consider measures of dispersion characterized by a single parameter. However, differences in dispersion may be more complex than can be characterized by a single parameter; for example, two groups can have very similar *mean* dispersion even while the *distribution* of pair-wise distances can be different. Such differences in dispersion would not be detected using a test that only compares mean square distances. Here, we develop the Distance-based Test of Homogeneity (`DTH`) that is sensitive to arbitrary differences in dispersion (as measured by the distribution of within-group distances) between multiple groups, or across values of a continuous variable.

The remainder of this work is organized as follows. In Section 2 we introduce two motivating examples; in Section 3, we describe `DTH` and the permutation scheme we use to assess significance. In Section 4 we describe simulations we undertook to compare `DTH` with the methods of Anderson and Gijbels and Omelka, concentrating on situations where differences in dispersion occur in the distribution of distances, rather than differences in single parameter. In Section 5 we analyze data on the oral microbiome of 511 participants by smoking status (never, former and current smokers) and by HIV status (HIV positive or negative). Finally, Section 6 contains concluding remarks. Codes and simulations to replicate the results provided in this paper, along with vignettes and datasets used in this paper can be found in https://github.com/asmita112358/DTH.

## 2 Motivating examples

We give three examples of situations where testing for dispersion is important. Two of them are cases where tests that are sensitive to more than mean square distances are important, the other is an example of ecological data where PERMANOVA is non significant, but tests for dispersion are.

The first example is from gut microbiome data on patients with inflammatory bowel disease (IBD) (Lloyd-Price et al. [2019]). Specifically, we consider baseline data from stool microbiome data among 32 participants diagnosed with Crohn’s Disease (CD) and 20 participants diagnosed with Ulcerative Colitis (UC). It is of interest to see how the gut microbiome differentiates these two disease categories that are typically diagnosed by clinical symptoms. In Figure 1, we show violin plots (of the log-transformed) within-group distances calculated using the Bray-Curtis distance, and also show the empirical distribution functions of the (untransformed) distances. Even on the log scale, the group medians are quite similar; however, there is clearly more dispersion in the CD group than the UC group. This is also seen in the two empirical distribution functions. In this example, we see that neither `betadisper`(p-value: 0.125) nor GO or GO.perm (p-values: 0.104 and 0.117 respectively) find a significant difference in dispersion, while `DTH` does (p-value: 0.024).

**Figure 1.**
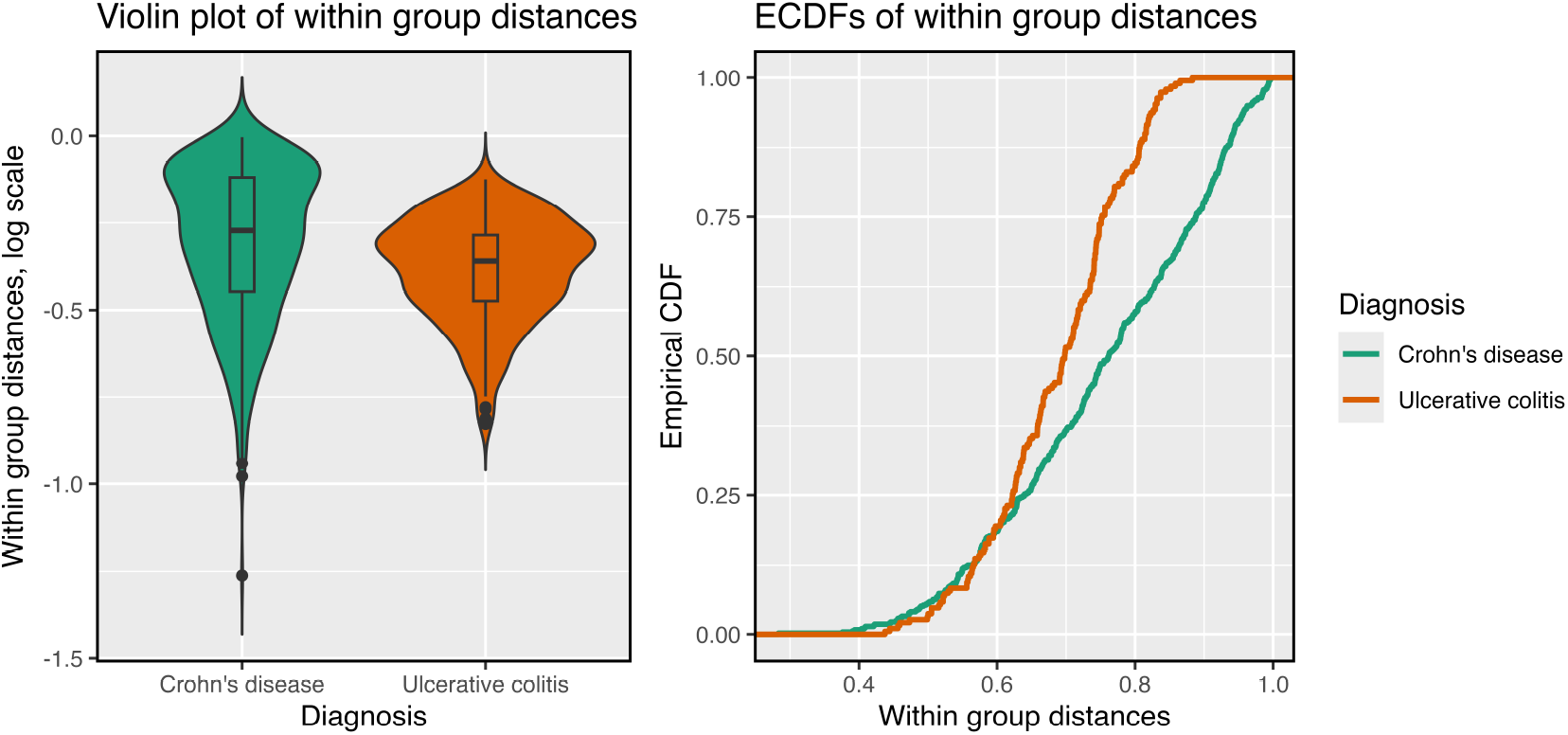
Violin plot and ECDF of the within-group distance for the two diagnosis groups in the IBD data as in the first motivating example

A second example arises when comparing methods for simulating synthetic microbiome data using the entire longitudinal IBD data as template, i.e the IBD data containing microbiome data from all visits of all patients. When developing the `MIDASim` (He et al. [2024]) method, we noticed that both `MIDASim` and the `SparseDossa` (Ma et al. [2021]) methods had mean square distances matching the template data they were trying to replicate. However, the `SparseDossa` data had a much narrower distribution of pairwise distances than that in the `MIDASim` data, which better matched the template data. Figure 2 offers a visual perspective of the problem, and table 1 reports the corresponding p-values.

**Table 1:**
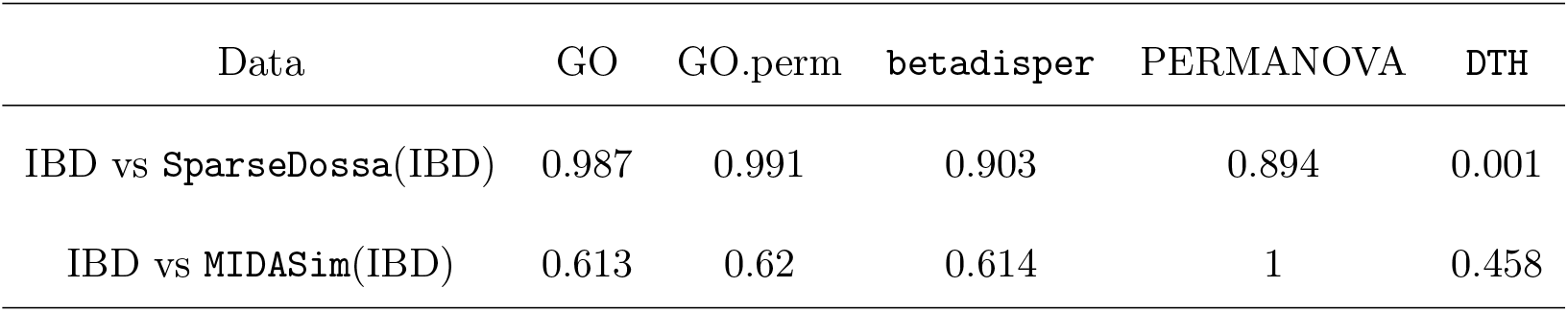
P-values testing dispersion homogeneity between true microbiome samples from a study of IBD and simulated datasets generated by SparseDOSSA2 and MIDASim using the true samples as templates.

**Table 2:** Scheme for permutation p-value computation. Here we have function values **m**_0_, **m**_1_, …, **m**_*R*_, each of which is a vector of length 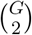 Note that p-values can be obtained for each column from a single set of permutations by using the replicates in the preceding columns as the null distribution for that quantity.

| Source | K | W | $M(\mathbf{p}^K, \mathbf{p}^W)$ | Pairwise p-values<br>$\mathbf{p} = (p_{12}, p_{13}, p_{23})$ | $F(p_{12}, p_{13}, p_{23})$ |
| --- | --- | --- | --- | --- | --- |
| Original data → | $\mathbf{p}_0^K$ | $\mathbf{p}_0^W$ | $\mathbf{m}_0$ | $\mathbf{p}_0$ | $f_0$ |
| Replicate 1 → | $\mathbf{p}_1^K$ | $\mathbf{p}_1^W$ | $\mathbf{m}_1$ | $\mathbf{p}_1$ | $f_1$ |
| Replicate 2 → | $\mathbf{p}_2^K$ | $\mathbf{p}_2^W$ | $\mathbf{m}_2$ | $\mathbf{p}_2$ | $f_2$ |
| ⋮ | ⋮ | ⋮ | ⋮ | ⋮ | ⋮ |
| Replicate R → | $\mathbf{p}_R^K$ | $\mathbf{p}_R^W$ | $\mathbf{m}_R$ | $\mathbf{p}_R$ | $f_R$ |

**Figure 2.**
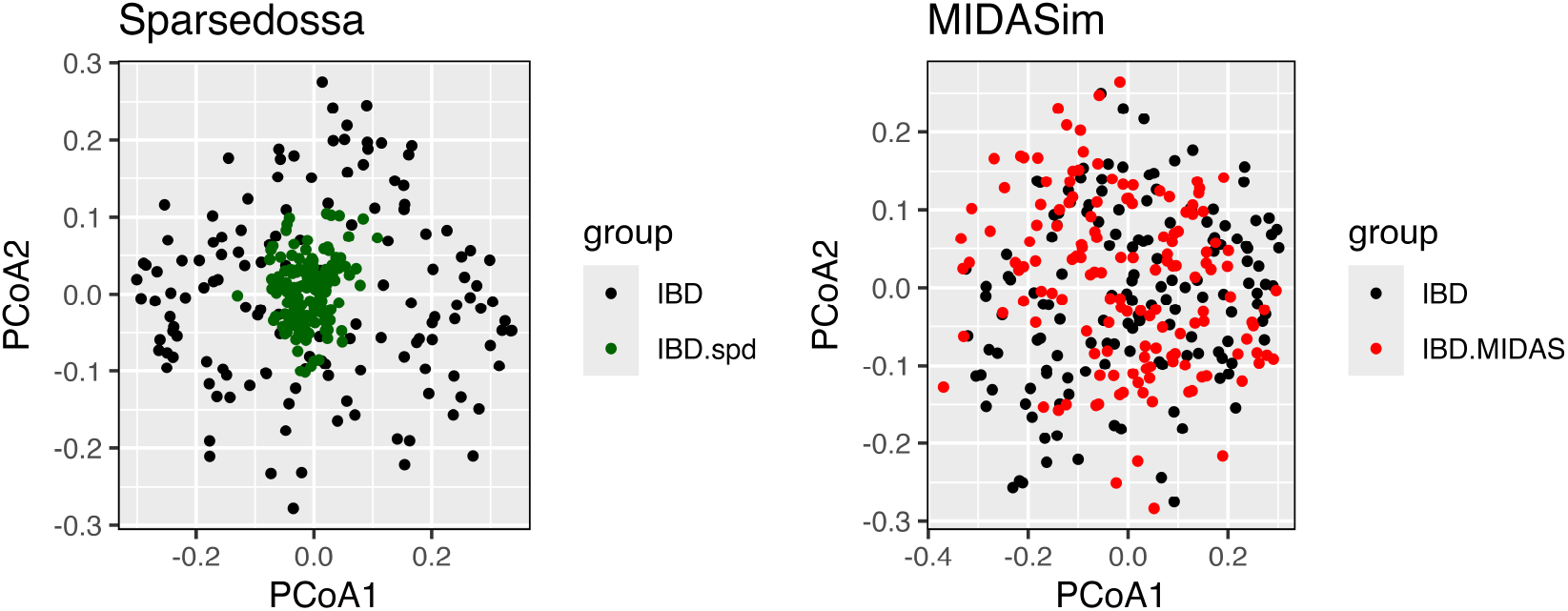
PCoA plots for Jaccard distance on the real IBD data (black) vs IBD as generated using `SparseDossa`(green, left panel) and `MIDASim`(red, right panel), respectively

In the third example, we consider Bumpus’ Sparrows Data (Bumpus [1899]) which consists of five morphological characteristics of sparrows measured in Rhode Island, grouped by survival status after a severe storm. We consider a subset of the data considered in Manly [2005] and Gijbels and Omelka [2013], and test for differences in dispersion amongst morphological characteristics by group. It is evident from Figure 3 that there is a significant difference in the dispersion of morphological characteristics between species that survived vs species that didn’t, and this is backed up by the p-values (in caption). The purpose of this example is to demonstrate the applicability of our proposed test beyond high dimensional data.

**Figure 3.**
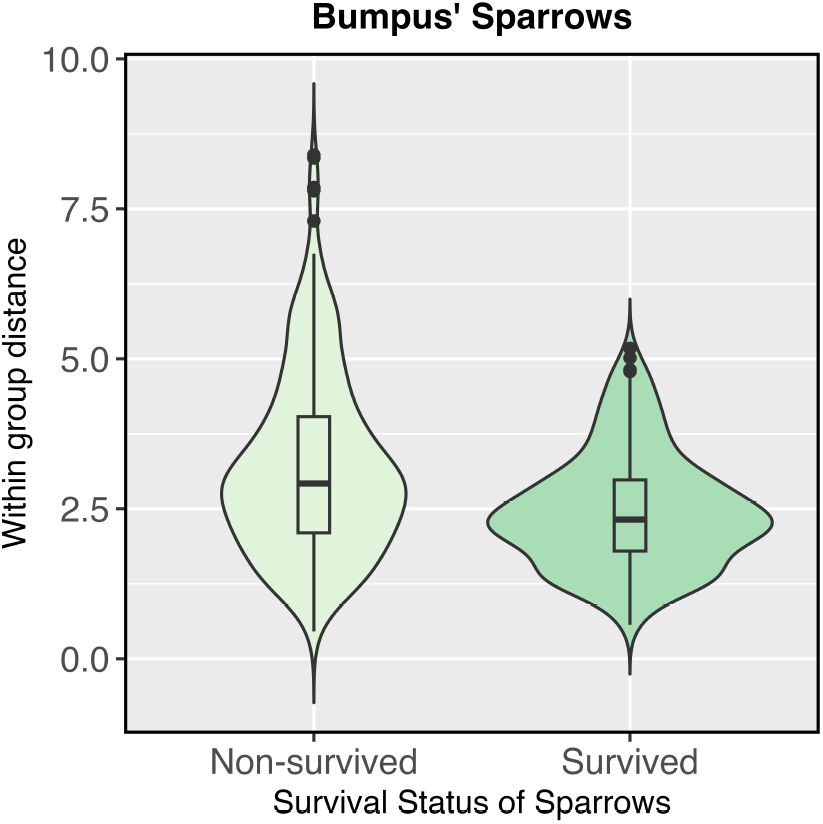
Distribution of within-group differences for the characteristics of species of sparrows that survived vs the ones that didn’t. pvalues: GO: 0.043, GO.perm: 0.057, PERMANOVA: 0.839, Betadisper: 0.065, `DTH`: 0.041

## 3 Methods

We assume that the data are summarized in a *n* × *d* matrix *Y* such that each row corresponds to an observation and each column to a taxon. We let *Y*_*i·*_ denote the *i*th row of *Y*. These samples are organized into *G* grou we let *g*_*i*_ denote the group that the *i*th observation belongs to. The *g*th group contains _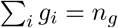_ samples and 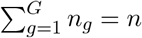. In this section, we will first consider scenario when we have samples from *G* = 2 groups. We will discuss how to extend this approach to three or more groups in Section 3.3.

For *G* = 2, let ℒ_1_ and ℒ_2_ denote the distributions of the within group distances for groups 1 and 2, respectively. We are interested in the following broad class of null hypothesis, namely:

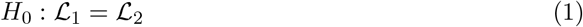

Define the distance between the *i*^*th*^ and *j*^*th*^ sample as

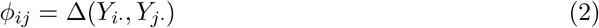

where Δ denotes the dissimilarity measure. This may be a Euclidean distance, or for microbiome data, it can be Bray-Curtis or Jaccard dissimilarity. Following Dubey et al. [2024] we can define for the *i*^*th*^ sample in the *g*^*th*^ group (*g* = 1, 2)

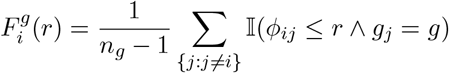

which is the proportion of samples in group *g* which is within distance *r* of the *i*^*th*^ observation.

The mean of this quantity, taken over samples in group *g*, is

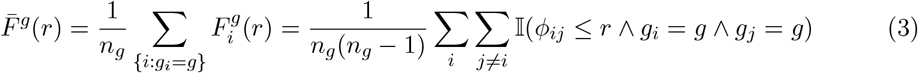

Considering this quantity as the building block of our test statistic instead of mean within group distance allows us to quantify the overall difference in distribution instead of the differences in mean dispersion only, hence paving the way for a broader class of hypothesis. Note that, 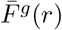 is a step function with at most 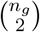 jumps. Testing for equality of ℒ_1_ and ℒ_2_ would translate to testing for equality of the functions *F*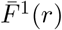 and *F*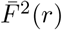. Two classical approaches of measuring this are by the Kolmogorov-Smirnov distance(Kolmogorov [1933]) or the 1-Wasserstein Distance (Vallender [1974], Kantorovich [1960]) between 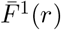 and 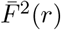.

It is known that the Kolmogorov-Smirnov test suffers from low power against all but location scale alternatives, (Zhou et al. [2017], Fan [1996]), while Wasserstein distance is more powerful against minor geometric differences (e.g outliers, differences in tail behavior) between the two distributions(Raghvendra et al. [2024]). In our motivating examples and applications, we encountered both types of differences between the groups, so it made more sense to combine the two types of distances, Kolmogorov-Smirnov and Wasserstein, and construct an omnibus test. This allows us to harness the best of both worlds, and results in a test that is powerful against both location-scale alternatives and differences in tail behavior.

### 3.1. DTH for Two Groups

For each group we compute 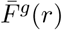, *g* = 1, 2, and measure the Kolmogorov-Smirnov (*K*) and 1-Wasserstein (*W*) distance between 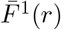 and 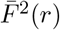. Since 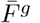 is a step function, both *W* and *K* can be easily calculated. We used the function `wasserstein1d` in the R package `transport` (Schuhmacher et al. [2020]) to calculate *W* the R function `ks.test` to calculate *K*. Because the within-group distances are not independent, asymptotic null distributions cannot be used to assess the significance of either *W* or *K*. To estimate the null distributions of *W*, *K* and their combinations we used the permutation approach described in Gijbels and Omelka [2013] to generate *R* replicate datasets in which the dispersion is the same in each group.

We denote by *K*_*r*_ (*W*_*r*_) the value of K (W) statistic for the *r*^*th*^ permutation replicate for *r* = 1 … *R*, and let *K*_0_ (*W*_0_) denote the value of K (W) for the original data. The *p*-value 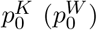 for the K (W) statistic is given by *p*^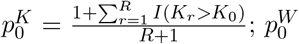^ is obtained by replacing *K* with *W*. Note that *p*-values for the null replicates can also be assigned using 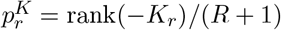and 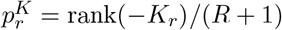.

We can combine the p-values from the K and W tests to form an omnibus test. Based on simulation results (not shown) we used *m* = 1 − min(*p*^*K*^, *p*^*W*^)) to combine the K and W tests, resulting in a statistic where larger values imply evidence against null. Using the *p*-values 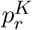 and 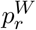we can calculate the value of 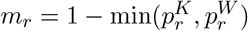for the original data (*r* = 0) and each null replicate (1 ≤ *r* ≤ *R*). When there are only two groups, we can then obtain 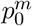, the *p*-value of the omnibus test, using 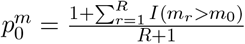

### 3.2 DTH for More than Two Groups

When we have more than *G >* 2 groups, we compute the omnibus statistic *m*_0;*gg′*_ for the original data and null replicates *m*_*r*;*gg′*_ for each pair of groups *g, g*^*′*^ as described in Section 3.1.

We then assign *p*-values 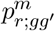_*′*_to each null replicate *m*_*r*;*gg′*_ using their rank among the null replicates as described previously. Based on simulations (results not shown) combine the 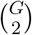 *p*-values 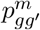 _*′*_ values using Fisher’s combination statistic 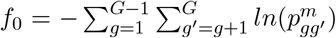 Values of *f*_*r*_ for each null replicate can be obtained by replacing 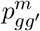 by 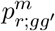 in the expression for *f*_0_. The final significance is based on 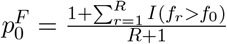 We note that we could have combined the 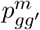 values using the Cauchy Combination test Liu and Xie [2020] which allows an analytic *p*-value to be assigned, but found the Fisher combination had better performance and only adds a trivial amount of computational effort.

### 3.3 Extension to Continuous Variables

Although the methods developed up to this point were designed for grouped data, they can be directly applied to test the association of changes in dispersion with continuous covariates. Since our interest is in hypothesis testing, not estimation, we can discretize any continuous covariates and then test for association between the covariate-defined groups and dispersion. In fact, this approach can also be used for the GO and GO.perm Gijbels and Omelka [2013] approaches. In the supplementary material we present an example simulation that uses a continuous variable.

## 4 Simulations

We conducted simulations to assess the performance of `DTH` in evaluating differences in dispersion and to compare it with existing tests of differential dispersion. First, to ensure precise control over dispersion parameters, we generated multivariate data from multiple parametric distributions for *G* groups. In the interest of fairness, we consider two scenarios in each setting, one that favors our method over others, and another that favors existing methods. Additionally, we performed simulations using MIDASim and sparseDOSSA, two recently-developed programs designed to generate realistic microbiome data. Both programs were designed to replicate key characteristics of real datasets, including high sparsity and large overdispersion. As shown in our motivating examples, these two methods can generate datasets with similar centroids but different dispersions. We also implement a simulation where the dispersion of the outcome is related to a continuous covariate, and report the results in the Supplementary materials.

### 4.1. Simulation 1: Multivariate Normal Distribution

We first simulated data from *d* -dimensional mixtures of normal distributions for *G*(*G* = 2, 3, 5) groups with identical group means but distinct distributions of dispersions. For each sample *i* in group *g*, we generate *Y*_*i·*_ | *v*_*i*_ ∼ *N*_*d*_(**0**_*d*_, *v*_*i*_**I**_*d*_), where *v*_*i*_ is generated from a mixture distribution, and the parameters of this prior varies from one group to another. We choose *d* = 500 for all simulations, and measured distance using the Euclidean distance.

We simulated four scenarios corresponding to four choices for the mixture distribution of *v*_*i*_ denoted by S0, S1, S2, and S3. These scenarios are ordered to grade progressively from scenarios that favor our method (S0, S1) to situations that favor existing tests (S2, S3). In S0 we have 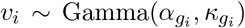 where 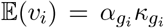. We choose the scale and shape 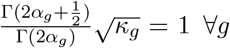 parameters such that *κ*_*g*_ = 1 ∀*g*; with this choice, *E*(*D*_*ii′*_)=Const when *g*_*i*_ = *g*_*i′*_ so that the GO tests and betadisper should have negligible power. In S1 and S2 we choose 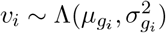 where Λ denotes the log normal distribution so that 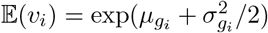 In S1 we choose 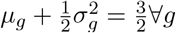 so that the mean dispersion 𝔼(*v*_*i*_) is constant across groups. Because the distribution of the dispersion differs between groups, this scenario favors our method, which is sensitive to differences in the distribution of dispersions. In S2 we keep *μ*_*g*_ constant ∀*g*, and vary 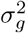 by group, so that the mean dispersion *E*(*v*_*i*_) also varies across groups. S2 favors existing methods that are designed to detect differences in mean dispersion. In S3, we choose a fixed value *v*_*g*_ for each group (see Table 3 for the values). We consider both balanced (B) scenarios, where each group has the same number of samples, and unbalanced scenarios (UB), where group sizes differ. The results for the balanced setting is presented in the main paper, and the unbalanced setting is presented in the Supplementary materials. The values of the mixing distribution parameters and sample sizes for each simulation are given in Table 3. Each parameter is expressed as a function of *θ*, a parameter which controls the deviation from the null hypothesis. In all cases, *θ* = 0 corresponds to the null hypothesis that the distribution of *Y* is identical in each group. Table 3 summarizes the simulation setup.

**Table 3:**
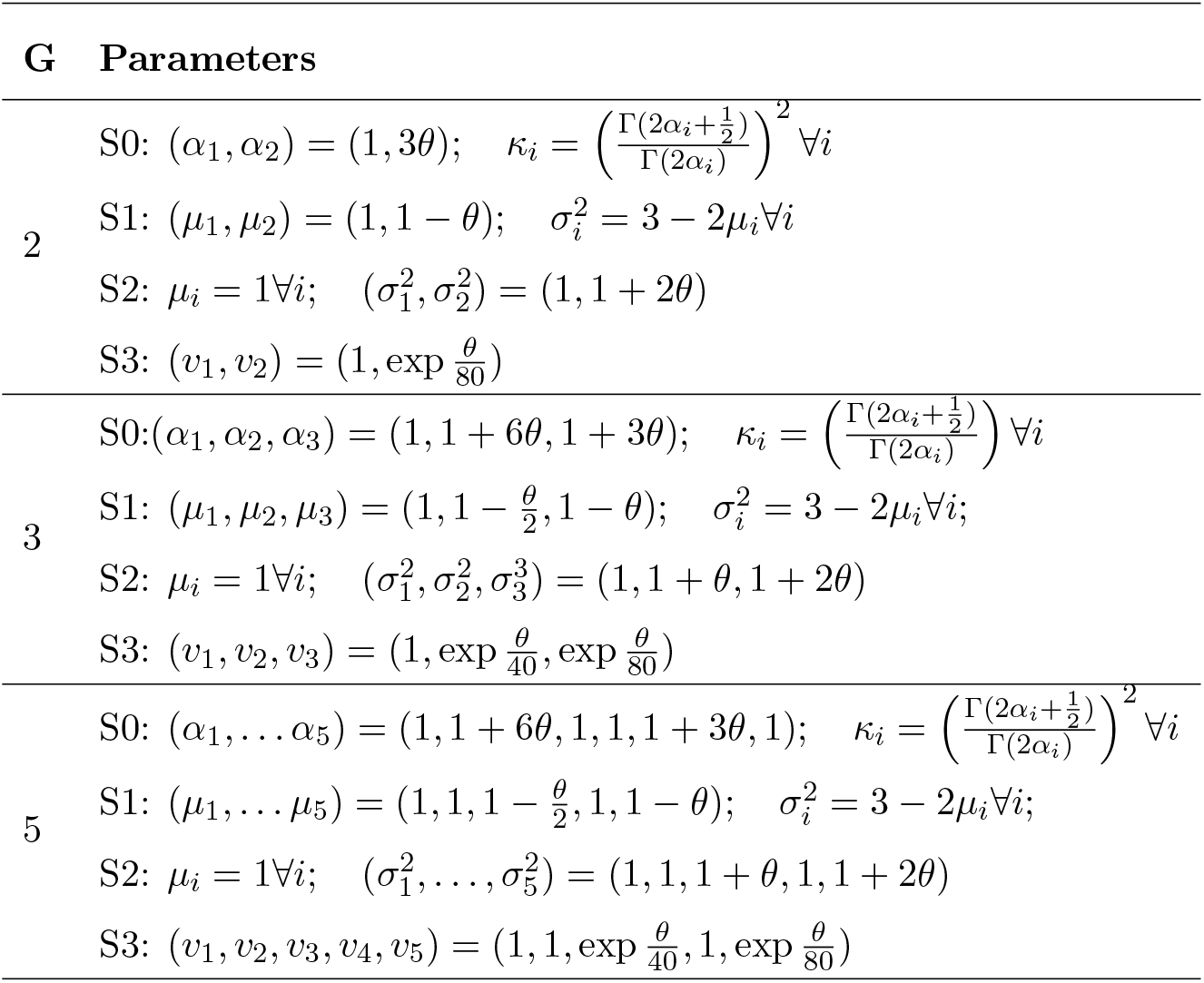
Simulation setups for multivariate normal outcome. *θ*, the degree of departure from null, varies from 0 to 2 in increments of 0.5 in every scenario. *d* = 500 and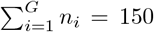 across all simulations.

In Figure 4 we show the empirical power for each test for each scenario and three values of *G*. At *θ* = 0 we see that each test controlled size. In every scenario but S3, `DTH` had noticeably higher power than the other tests. Even in S3, `DTH` had comparable or slightly higher power than the other tests. `PERMANOVA` had essentially zero power. `Betadisper` performed second best in S0 and S1, while the GO tests were second best in S2.

**Figure 4.**
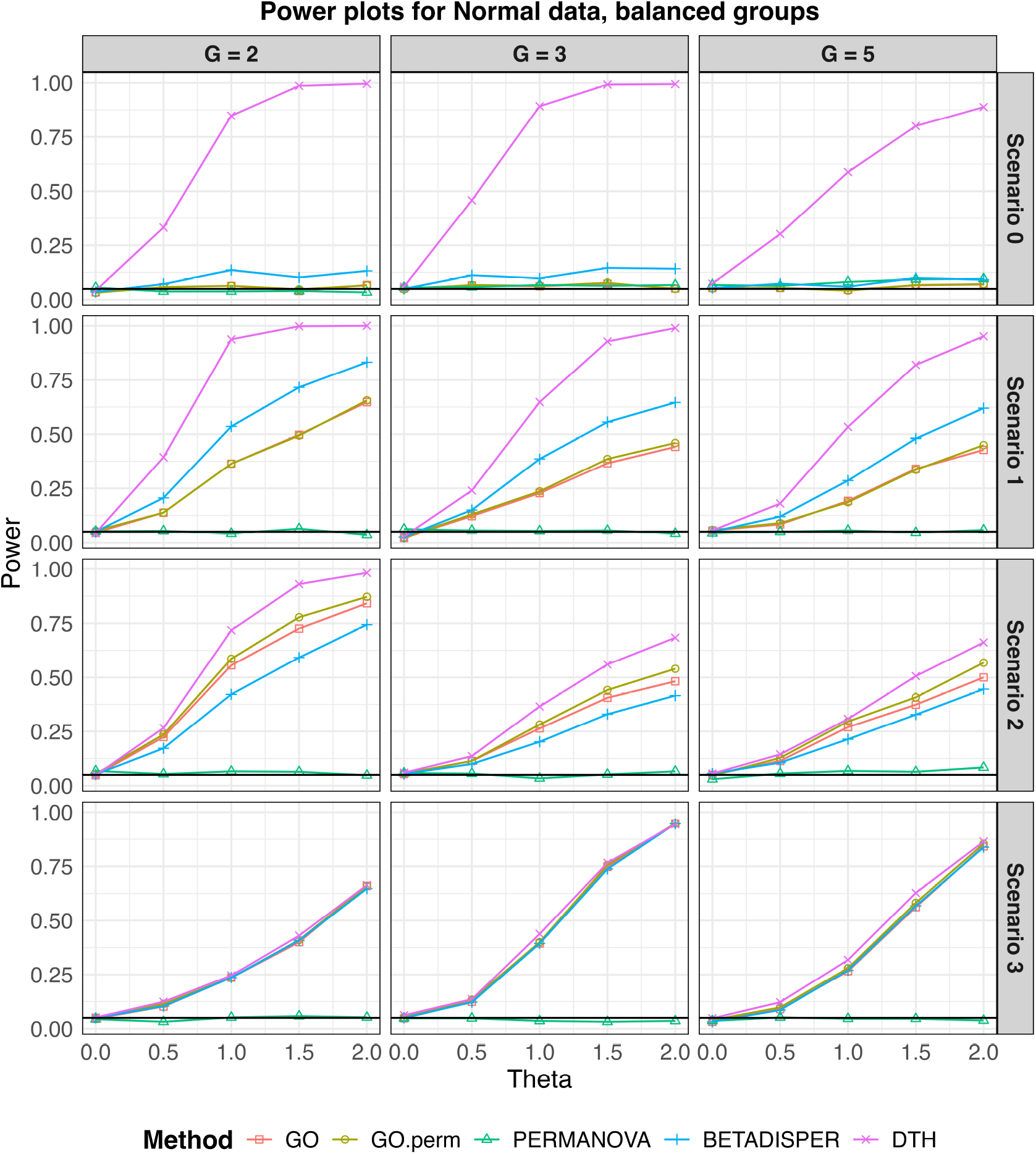
Empirical type I error and power for multivariate normal outcomes for balanced designs. Sample size for all simulations was 150 observations split evenly among the *G* (*G* = 2, 3, 5) groups. *θ* = 0 corresponds to the null scenario. A solid black line marks the nominal detection level (0.05).

**Figure 5.**
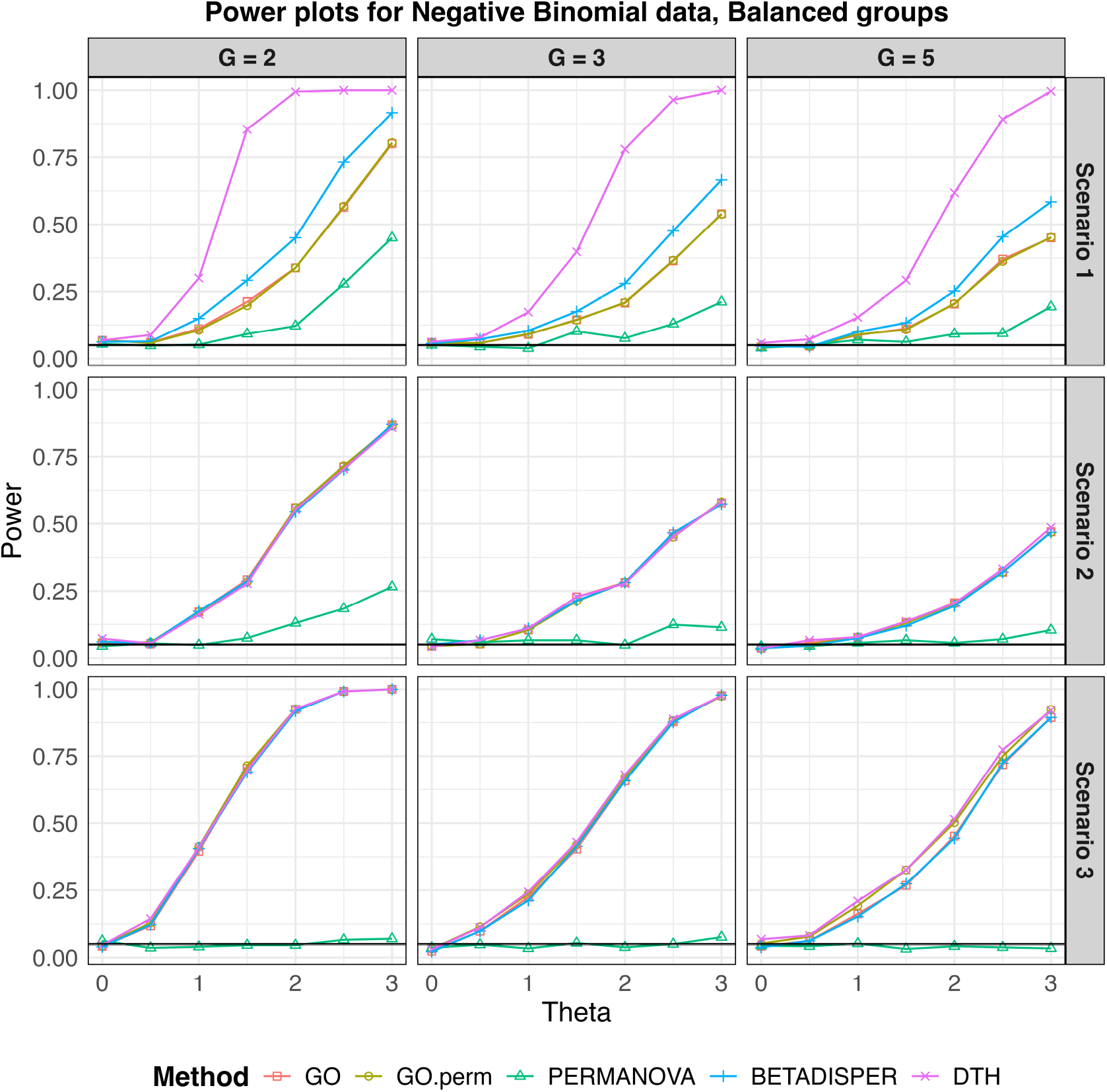
Empirical type I error and power for Negative Binomial outcome with balanced designs. Sample size for all simulations was 150 observations split evenly among the *G* (*G* = 2, 3, 5) groups. *θ* = 0 corresponds to the null scenario. A solid black line marks the nominal detection level (0.05).

### 4.2. Simulation 2: Negative Binomial Distribution

Because microbiome data are often count data, we generated data *Y*_*ij*_ from a mixture of negative binomial distributions. For observation *i*, we generated data for 500 taxa using 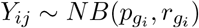 where 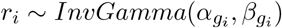. We chose NB parameter *p*_*g*_ = *r*_*g*_*/*(*μ*_*g*_ +*r*_*g*_) so that 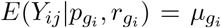 and 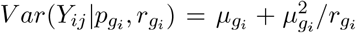 hence the unconditional moments of *Y* are *E*(*Y*_*ij*_) = *μ*_*gi*_ and 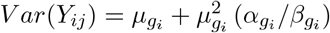 Here, we consider three scenarios S1-S3 that are comparable to S1-S3 for the normal simulations in 4.1. We do not include a scenario analogous to S0 as it is difficult to construct this case. Distance was measured using the Bray-Curtis distance.

For S1, we vary the parameters *α*_*g*_ and *β*_*g*_ across groups such that 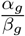 remains constant. This ensures that the mean dispersion remains the same in all groups but the distribution of the dispersion varies. For S2, we held *α*_*g*_ constant across groups while varying *β*_*g*_. This creates a simulation setting that favors the existing method by changing mean dispersion, i.e 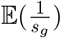 across groups. For S3, *r*_*i*_, the dispersion parameter associated with the Negative Binomial Distribution is deterministic, and changes across groups in the non null settings. The number of groups, their sizes, and parameter values as a function of a parameter *θ* are given in Table 4. For each scenario, we consider a balanced (B) and unbalanced (UB) setting, where the results for the balanced setting is presented in the main paper, and the unbalanced setting is presented in the Supplementary materials.

**Table 4:**
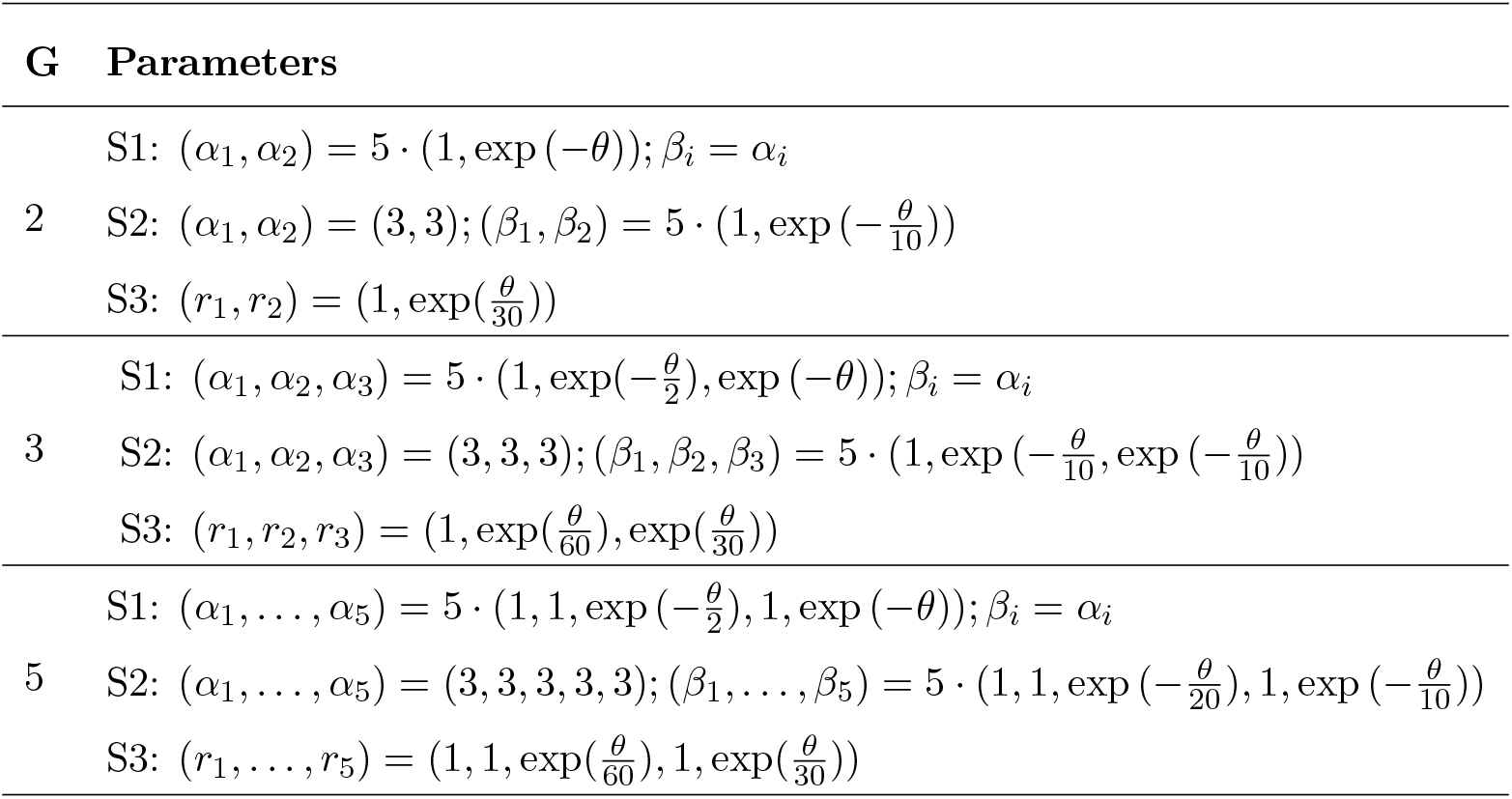
Simulation setups for Negative Binomial outcome. *θ*, the degree of departure from null, varies from 0 to 3 in increments of 0.5. *d* = 500 and 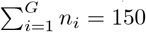 across all simulations.

We find that all methods have similar empirical size. For the alternative model, in S1, `DTH` outperforms the competing methods significantly, and in S2, it has comparable power to GO, GO.perm and Betadisper.

### 4.3 MOMS-PI data

To evaluate the performance of `DTH` on realistic microbiome data, we compared data from the Multi-Omic Microbiome Study–Pregnancy Initiative (MOMS-PI) Fettweis et al. [2019] with simulated data generated using sparseDOSSA Ma et al. [2021], which was designed to mimic the characteristics of the MOMS-PI dataset. The original MOMS-PI and simulated data each have 514 samples with 1146 taxa. Given this large sample size, all methods yield highly significant p-values (<0.001). To generate meaningful comparisons, we randomly sampled between 50 and 250 pairs of observations from the two datasets, and computed p-values for each subsample. The reported power is the proportion of subsamples that showed a significant difference (at the p=0.05 level) out of 1000 subsamples. We used all 1146 taxa in each subsample, and all competing methods were evaluated using both the Jaccard and Bray Curtis dissimilarities. The resulting power values are reported in Figure 6. We see that `DTH` has higher power to detect the difference between the real and simulated data at much smaller sample sizes than the other methods.

**Figure 6.**
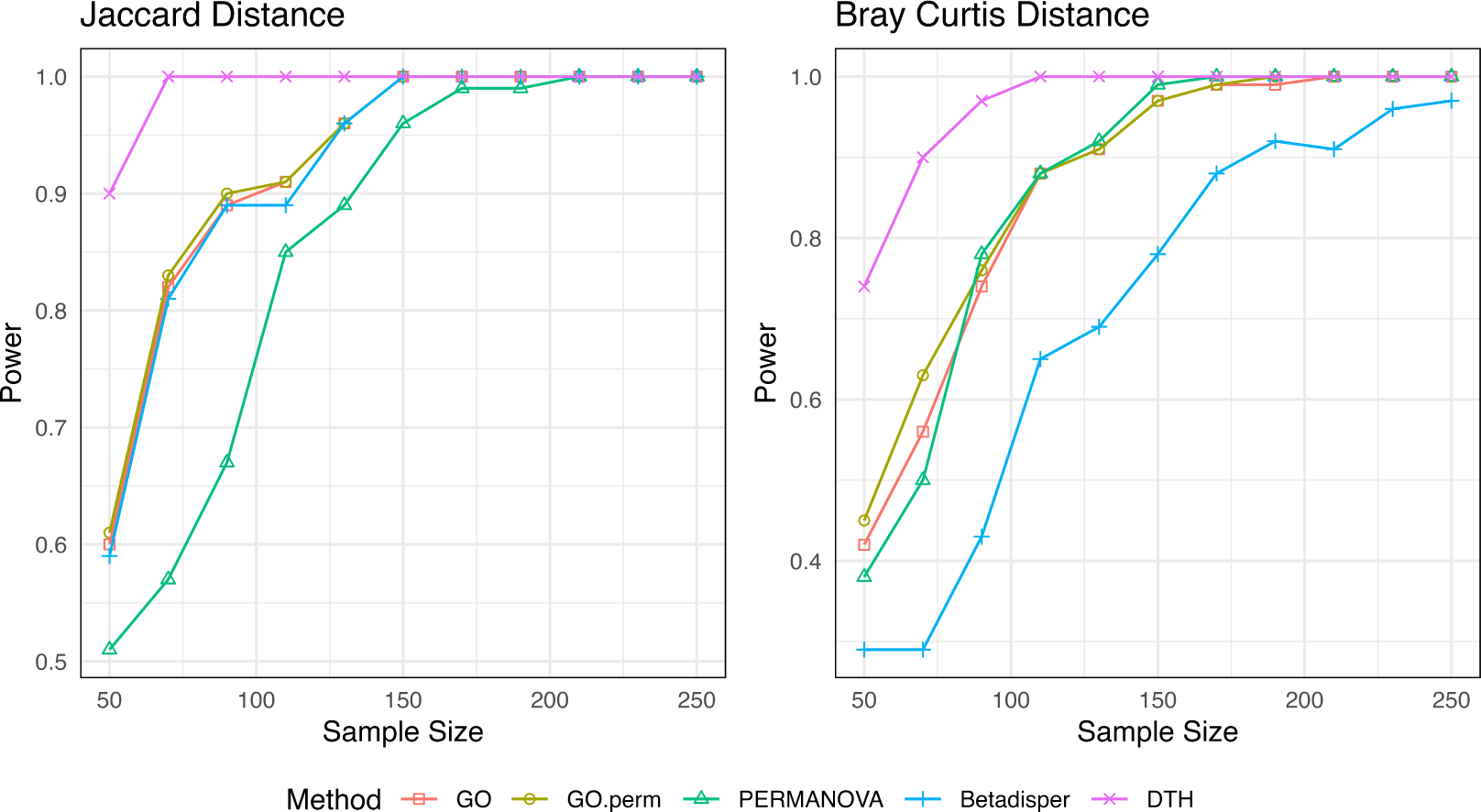
Empirical power of all methods based on 1000 randomly selected subsamples at smaller sample sizes(50 to 250). This simulation is intended for comparing dispersion of MOMS-PI data and Sparsedossa-simulated version of MOMS-PI at a smaller scale, and observing which method starts catching the differences in dispersion more effectively at lower sample sizes.

### 4.4 IBD data with MIDASim

To evaluate the performance of `DTH` on microbiome data under a controlled simulation setting, we based our simulations on the IBD dataset described in Section 2, which includes 32 participants with Crohn’s Disease (CD) and 20 with Ulcerative Colitis (UC). We first removed taxa that were entirely absent in either group, resulting in 327 taxa retained for analysis. For each simulation replicate, we used MIDASim to generate 52 CD-like microbiome profiles, using the original CD participants as the template, and 20 UC-like profiles, using the UC participants as the template. When generating simulated CD data, we preserved the original 52 observed library sizes from the CD group.

To study the power of the competing methods, we generated data that smoothly transitions from CD-like to UC-like. To accomplish this, we introduced a parameter *θ* such that, with probability 1 − *θ* we would replace the *i*^th^ UC microbiome profile with the 32 + *i*^th^ CD microbiome profile. This replacement was independently decided for each UC observation.

Thus, when *θ* = 0 all data are generated from the CD template, while when *θ* = 1 all CD samples are generated from the CD template and all UC samples are generated from the UC template.

The results of these simulations is shown in Figure 7. `PERMANOVA` has the highest power; this is presumably explained by noting a location shift between CD and UC data, illustrated by a PCoA plot of a typical simulation replicate for *θ* = 1 (Figure 7, panel B). The remaining (pure dispersion) methods all have about the same power, with betadisper performing slightly better than `DTH` but both GO method performing worse. This indicates that even in complex microbiome data, `DTH` has comparable power to GO and betadisper even though it tests a much broader alternative.

**Figure 7.**
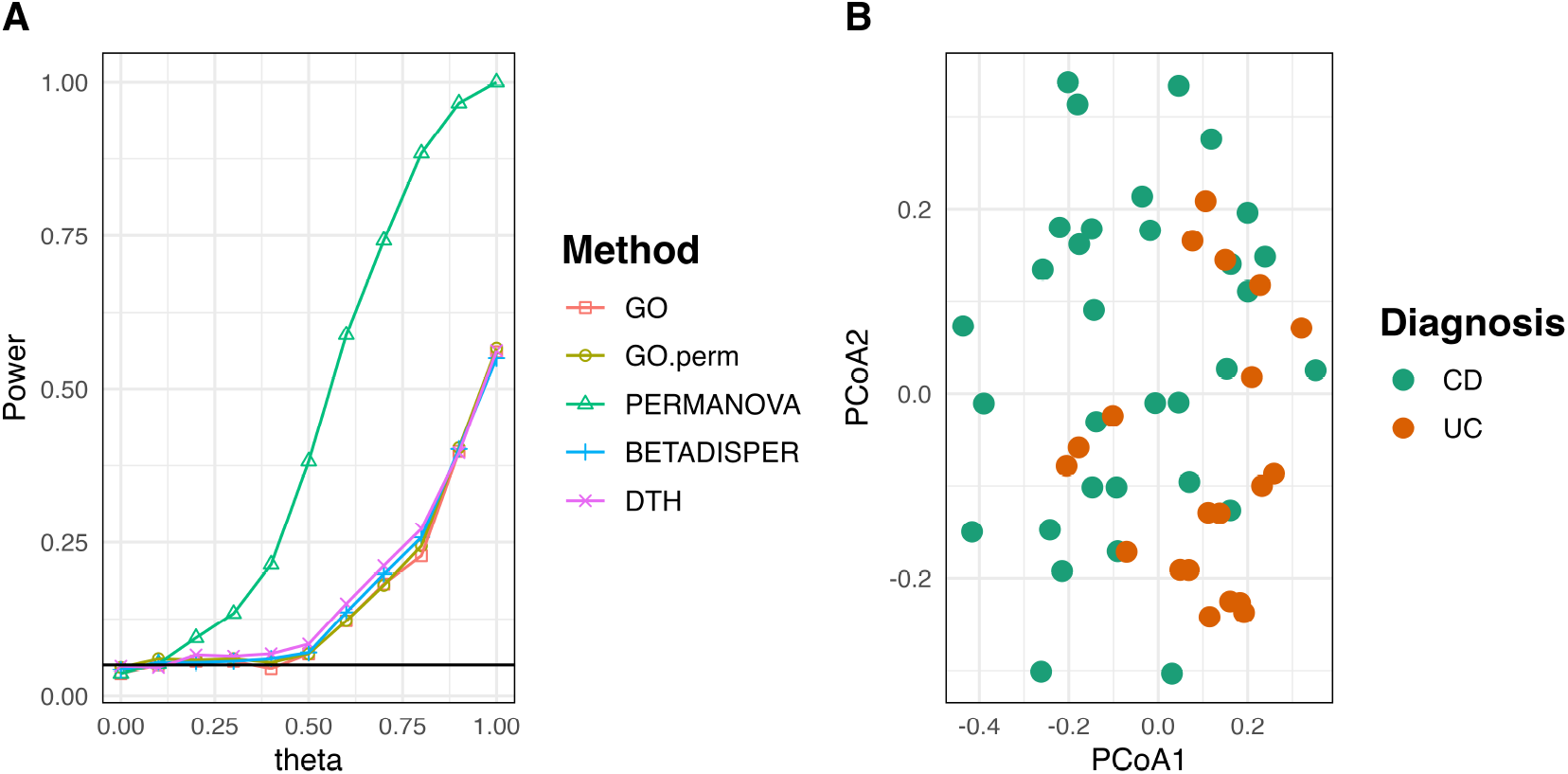
A) Empirical rejection rates for all competing methods for *θ* ranging from 0 to 1. *θ* = 0 corresponds to the null scenario. B) PCoA plot for CD vs UC cohort of synthetic IBD data generated using MIDASim

## 5 Analyses of Oral Microbiome Data

In addition to the data we’ve already analyzed in the motivating examples, we also analyzed data from the Men and Women Offering Understanding of Throat HPV (MOUTH) study Zhang et al. [2022]. Using data from this study, we compared the dispersion in oral rinse microbiome profiles between smokers and non-smokers, and persons with and without HIV infection.

For the HIV analysis, data from 524 participants were used, where 443 (84.5%) tested HIV-negative and 81 (15.5%) tested HIV-positive. For the smoking analysis, data from 511 participants were used and grouped by smoking status: never smokers (281 participants), former smokers (150 participants), and current smokers (60 participants). Violin plots depicting the distributions of the within-group distances are provided in Figure 8. The tests for differences in dispersion across groups (Table 5) yielded significant *p*-values for all competing methods (*p <* 0.005). Thus, these analyses serve as examples where `DTH` performs at least as well as the other tests. Interestingly, although the mean between-observation Bray Curtis distances of the groups are similar (0.51, 0.53, 0.55) for the smoking analysis and (0.52,0.58) for the HIV analysis, GO, GO.perm, and `betadisper` are as successful in finding differences in dispersion as `DTH`.

**Table 5:**
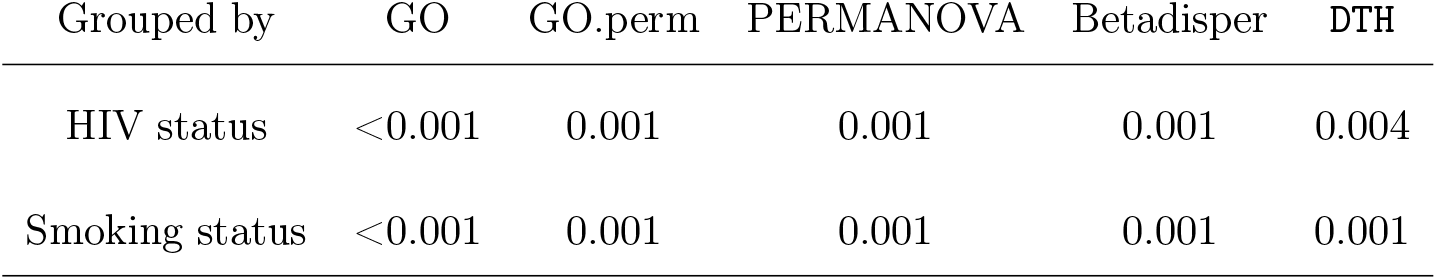
p-values for test of homogeneity of oral microbiome data across cohorts grouped by HIV status and Smoking status.

**Figure 8.**
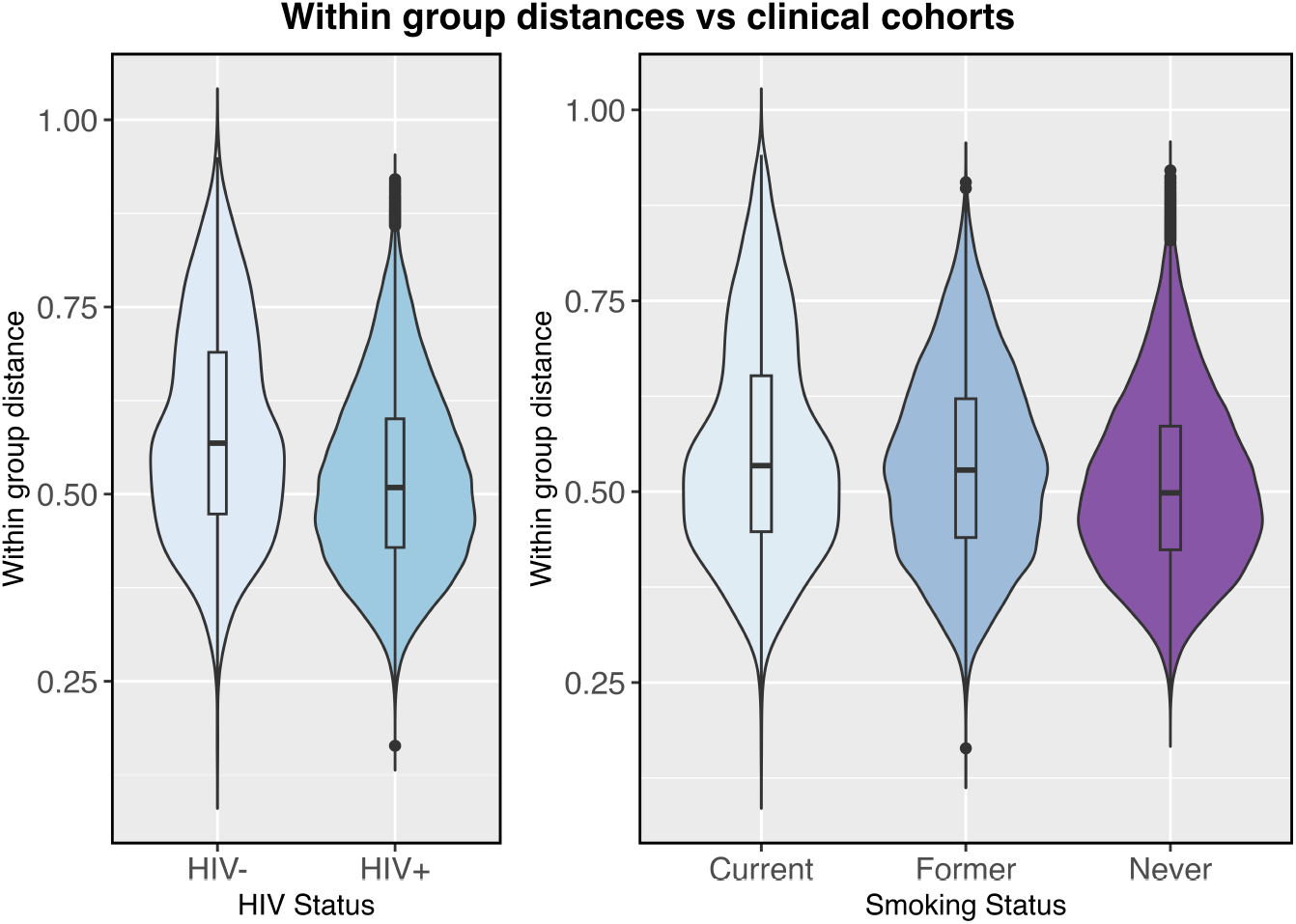
Distribution of within group distances of microbiome data vs different clinical cohorts (left: HIV status, right: smoking status) for MOUTH data

## 6 Discussion

We proposed a non-parametric test for dispersion that allows for more general alternative hypotheses than either of the two existing tests, Betadisper and the GO tests of Gijbels and Omelka [2013]. Our simulations and analyses of microbiome data indicate clearly that dispersion is more complex than can be captured by a single quantity. Dispersion can act at different scales; as our simulations show, simple tests may have very different performance when 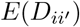 is similar across groups than when 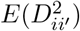 is similar. For this reason, a test that is sensitive to broad alternatives such as `DTH` is favored. Furthermore, our simulations show that, at least in the cases we considered, the loss of power incurred by using such a flexible test is, at worst, small.

We have illustrated the use of `DTH` in a number of applications, including assessment of the performance of large-scale omics data simulators. Dispersion tests can have inherent value. For example, we may find that case probands may exhibit higher diversity than controls (which may indicate unrecognized disease subtypes) or lower diversity (which may indicate a specific risk profile). Further, testing for differential dispersion is an important step in validating the assumptions of ANOVA and `PERMANOVA`, where it can be useful to verify the homoscedasticity assumption. However, the low power of `PERMANOVA` to distinguish pure dispersion alternatives in our simulations suggests that concern about `PERMANOVA` confusing dispersion and location may be somewhat overstated, at least when group sizes are balanced (`PERMANOVA` had non-negligible power for unbalanced groups; see Supplementary materials).

Although the emphasis and examples in this paper have all come from microbiome and ecological studies, this is not the only situation where we may be interested in dispersion across groups. Many types of non-numeric data can be converted to pairwise distances, such as forensic databases of fingerprint similarities. For data of this kind, tests of dispersion could be informative; for fingerprint data, we may want to see if dispersion varies across ethnic groups. Our tests also apply to numeric but high-dimensional data that can be converted to distances; for example, functional data could be converted into pairwise distances, after which distance-based approaches such as `DTH` could be used for further analysis. Although some features of the functional data may be lost, the gain in simplicity may be worthwhile.

## Supplementary materials

In the supplement, we present the additional simulation results for unbalanced groups for Normal and Negative Binomial data, as well as continuous covariates.

### Additional simulation results: Power for Unbalanced Groups

Figures 9 and 10 recapitulate results in Figures 4 and 5 but for unbalanced group sizes. See caption of figures 9 and 10 for group sizes. The results are similar; the only surprising feature is that PERMANOVA now has non-negligible power for Scenario 2 in both normal and negative binomial simulations.

**Figure 9.**
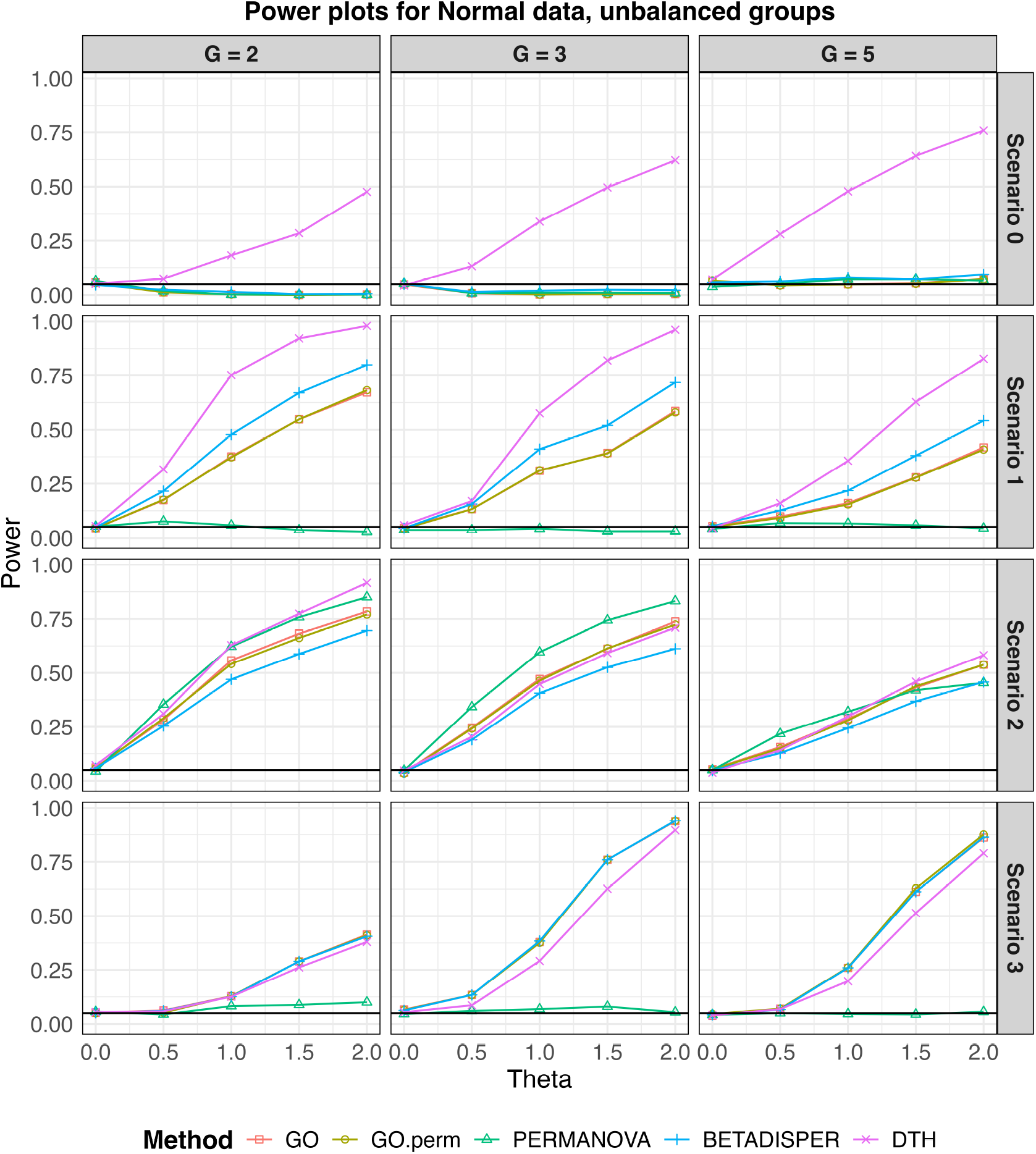
Empirical type I error and power for Multivariate Normal data with unbalanced designs (*G* = 2, 3, 5). The total sample size was fixed at 150. For two-group comparisons, sample sizes were 125 and 25; for three groups, 100, 30, and 20; and for five groups, 50, 40, 30, 20, and 10. *θ* = 0 corresponds to the null scenario.

**Figure 10.**
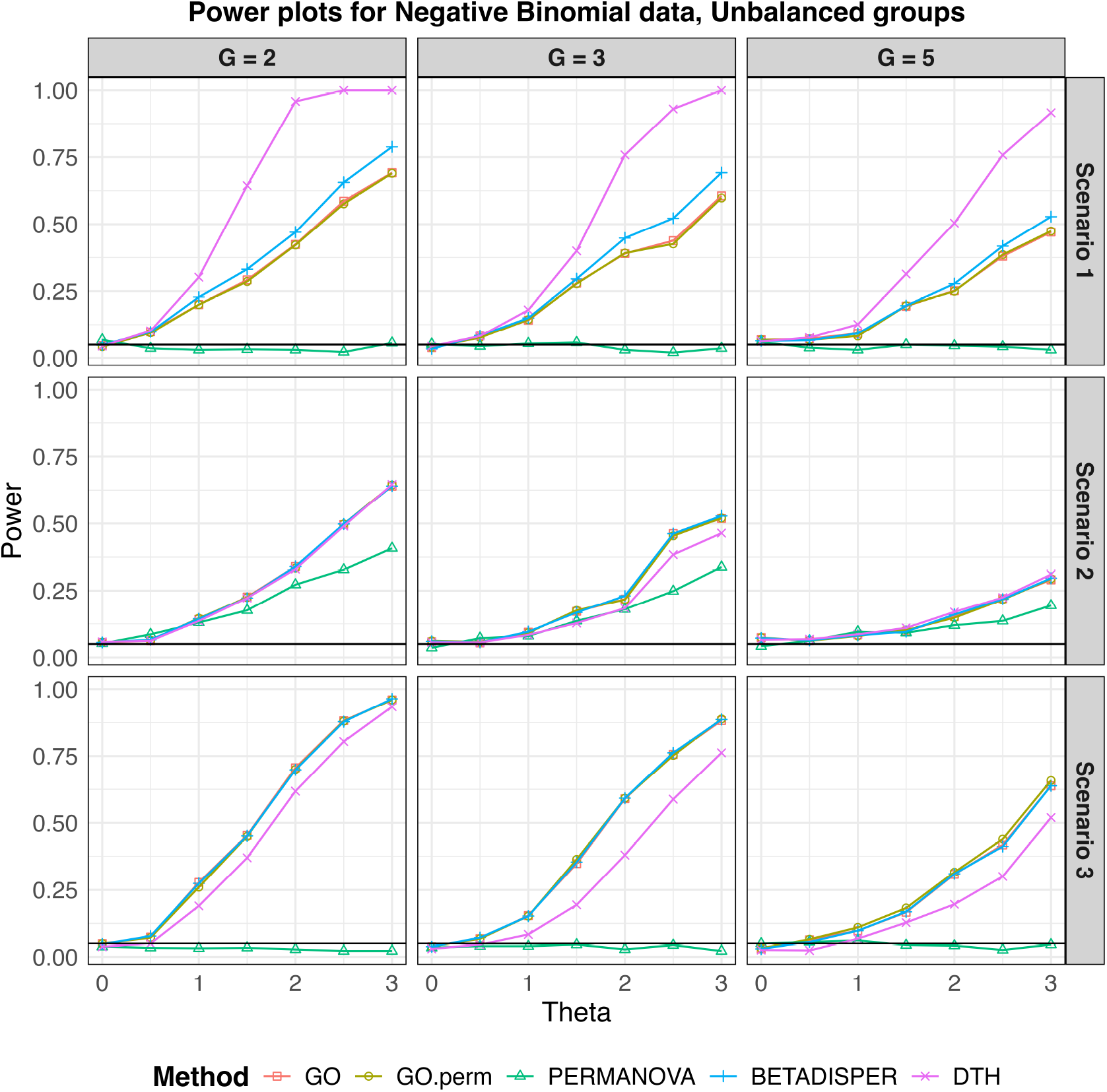
Empirical type I error and power for Negative Binomial data with unbalanced designs,(G = 2,3,5). The total sample size was fixed at 150. For two-group comparisons, sample sizes were 125 and 25; for three groups, 100, 30, and 20; and for five groups, 50, 40, 30, 20, and 10.

### Simulation Example with Continuous Covariates

As described in section 3.3, `DTH` can also be used to test for homogeneity with respect to continuous covariates. Here we demonstrate this using a simulation where heteroscedasticity is explained by a covariate *x*_*i*_ ∼ *Unif* (0, 5). We generate *n* = 150 observations *Y*_*i·*_, each having *d* = 500 components, using the model

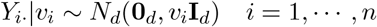

where

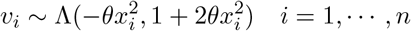

and where Λ(*µ, σ*) is the log-normal distribution. This choice ensures that 𝔼(*v*_*i*_) = exp (0.5) independent of covariate *x*, while the distribution of *v* does depend on the covariate; thus this simulation is similar to S1 in section 4.1. The parameter *θ* controls the degree of heteroscedasticity; *θ* = 0 corresponds to the null hypothesis, i.e no change in dispersion across values of *x*. To test for differences in dispersion across values of *x*, we bin *x* into intervals of length 1, and treat each bin as a group. Power for each tests considered here, for *θ* varying between 0 to 0.15, is reported in Figure 11. We observe that `DTH` is the most powerful amongst all competing methods.

**Figure 11.**
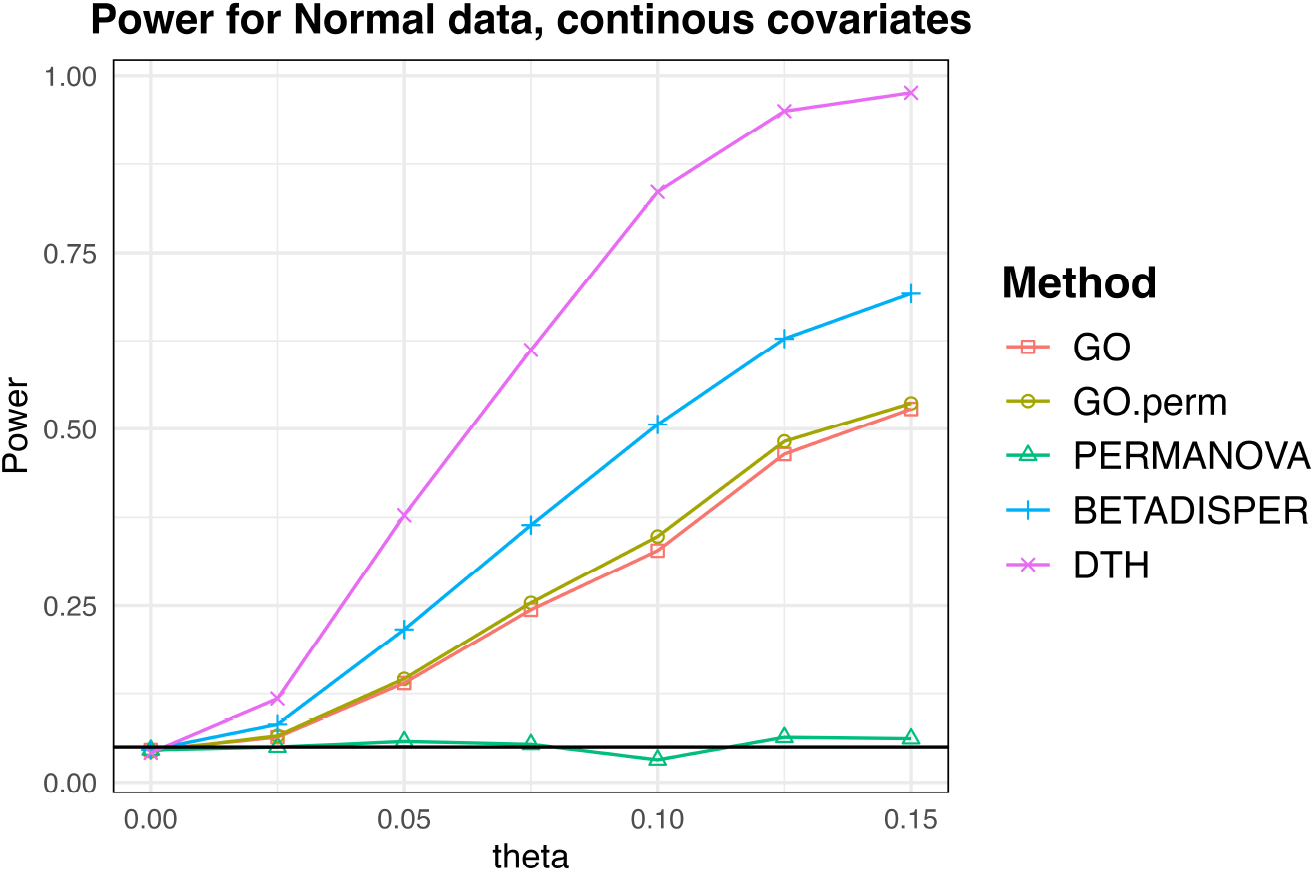
Empirical type I error and power for Multivariate Normal outcomes with continuous covariate. Nominal detection level (0.05) is marked with a solid black line.

